# Ribosomal Epistatic Interactions Drive the Emergence of High-Level Streptomycin Resistance in *Escherichia coli*

**DOI:** 10.64898/2025.12.13.694165

**Authors:** Fengjun Xu, Yue Xing, Yujie Men

## Abstract

The widespread use of antibiotics has raised serious concerns about the emergence and dissemination of antibiotic resistance in the environment. Studies reveal that environmental exposures to antibiotics and other non-antibiotic pollutants can promote the development of high-level antibiotic resistance via *de novo* mutations. However, specific genotypes conferring strong resistance in resistant mutants, particularly the roles of co-occurring mutations, remained poorly understood. This study fills this knowledge gap by constructing site-specific *Escherichia coli* mutants and demonstrating that it was the epistatic interactions that conferred the observed strong streptomycin resistance in native *E. coli* isolates. Three ribosomal mutations (*rpsE^D142N^*, *rpsL^R86S^* and *rsmG^W150fs^*), which co-evolved under environmentally relevant selection pressures, acted synergistically to confer resistance far exceeding that of each individual mutation. The interactions between the *rsmG* frameshift mutation and other reported *rpsL* point mutations were also investigated. The findings unravel the essential but previously overlooked role played by specific co-evolving mutations in the acquisition of both high-level streptomycin resistance and fitness compensation. The findings also underscore a strong and increasing need to investigate whole genomes of antibiotic-resistant bacteria and identify potential epistatic mutations conferring resistance phenotypes. The integration of those genetic mutations as antibiotic resistance biomarkers will complement with the resistome profiling and enable more accurate antibiotic resistance monitoring and environmental risk assessment.

**Synopsis:** Multiple ribosomal mutations co-evolved under environmentally relevant exposure conferred high-level streptomycin resistance via epistatic interactions.

Table of Contents Graphic

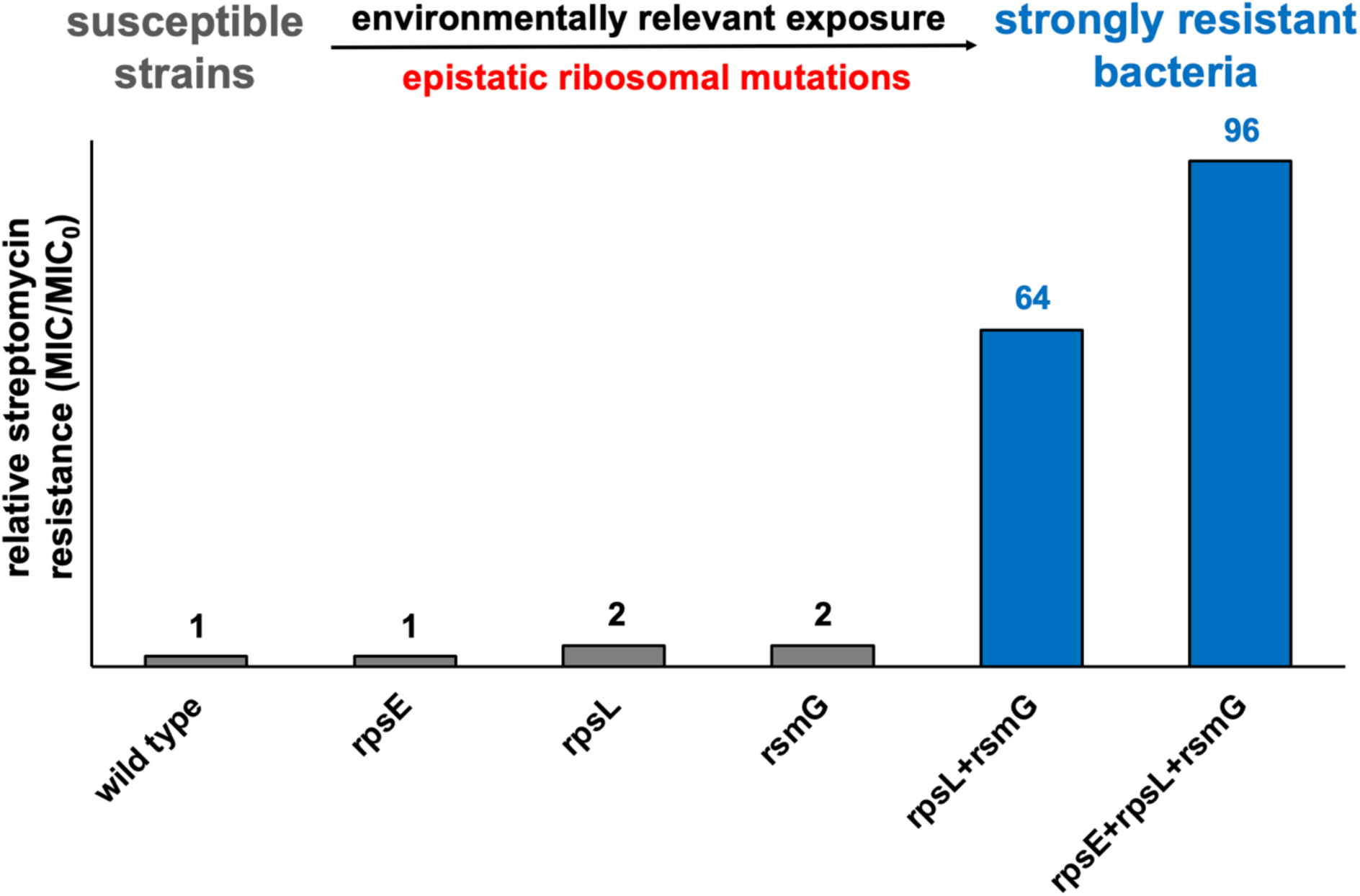

## Introduction

The widespread use of antibiotics has led to the emergence and dissemination of antibiotic resistance in the environment, posing one of the greatest threats to the health of humans, animals, and the environment.^1–3^ To mitigate the increasing risk of antibiotic resistance and promote One Health, substantial efforts have been made to control antibiotic usage and monitor antibiotic resistance determinants (i.e., antibiotics, antibiotic resistance genes, and antibiotic-resistant bacteria) in various environments. High-throughput sequencing-based antibiotic resistome profiling has become a widely applied tool to evaluate antibiotic resistance risks in a given environment.^4, 5^ However, resistome-based monitoring may involve false positives (i.e., genotypes do not necessarily lead to resistance phenotypes), especially for antibiotic resistance mechanisms involving target-altering mutations that reduce the antibiotic binding affinity to the target enzyme.^5^ Thus, it is crucial to identify and experimentally validate specific target mutations conferring antibiotic resistance, which may serve as complementary biomarkers or be integrated into existing antibiotic resistance gene databases for the surveillance of antibiotic resistance.

Resistance to several classes of antibiotics may be attributed to target alterations, including aminoglycosides (inhibiting ribosomal proteins). Among them, streptomycin is a commonly used one to treat bacterial infections in humans and animals.^6, 7^ It irreversibly binds to the 30S ribosomal subunit and disrupts protein synthesis. Bacteria can develop resistance by mutating genes encoding the target enzyme,^8, 9^ and the effectiveness of streptomycin is thus reduced due to the weakened drug-ribosome interactions. High-level streptomycin resistance is most frequently associated with mutations in *rpsL*, which encodes the crucial ribosomal protein S12 in the 30S ribosomal subunit.^10^ Loss-of-function mutations at *rsmG* (also known as *gidB*), which encodes the ribosomal RNA small subunit methyltransferase G, have been detected separately or together with *rpsL* mutations, and individual *rsmG* mutations may cause mild streptomycin resistance.^11, 12^ Those mutations were not only observed under clinically relevant conditions but also environments with low-level exposures to antibiotics and other non-antibiotic co-occurring pollutants. Our previous study revealed that co-exposure to low-concentration pesticides and streptomycin at environmentally relevant levels promoted the development of high-level streptomycin resistance in *E. coli* populations, where both target mutations (e.g., *rpsL^R86S^* and *rsmG^W150fs^*) were detected at the same frequency.^13–15^

Although individual target-altering mutations have been identified, evidence linking specific genotypes to the resistance phenotype is limited because in many previous studies only mutations of target genes were examined using Sanger sequencing, whereas the whole-genome information was missing. Consequently, there is a big knowledge gap regarding the interactions of co-evolved mutations in the genome and how they could contribute to resistance phenotypes. To date, there are only a couple of studies that incidentally touched those questions while aiming at other topics. One study showed that individual *rpsL* or *rsmG* mutations could act synergistically with the streptomycin-degrading gene (*strB*) on a plasmid, resulting in resistance levels much higher than those conferred by *strB* alone.^16^ However, the interactions between target-altering mutations at *rpsL* and *rsmG* in bacteria without resistant plasmids still remained unknown. In another study focusing on streptomycin production in *Streptomycetes* species, some results reflected a synergistic or additive effect on resistance in mutants, where both *rpsL* and *rsmG* mutations were detected by Sanger sequencing.^17^ However, since those mutants might harbor additional uncharacterized mutations not verified by whole genome sequencing, no concrete conclusion on epistatic interactions could be made. Therefore, a more specific and targeted investigation using appropriate molecular tools is urgently needed to interrogate potential epistatic interactions among co-evolved mutations that may have been previously overlooked but can shape high-level antibiotic resistance profiles.

Here, we systematically investigated the interactions among ribosomal mutations in *rpsL* and *rsmG* previously reported in streptomycin-resistant *E. coli* isolates. Site-specific mutants carrying individual or combined ribosomal mutations were constructed by introducing desired mutations into the chromosome of a wild-type, streptomycin-susceptible *E. coli* strain. The resistance level of each constructed mutant was compared. An epistatic effect of specific ribosomal mutations on the development of high-level resistance was observed and corroborated by reversing the mutations to the wild-type allele in the native resistant strain. The implications of those new findings for future antibiotic resistance studies, environmental monitoring, and risk assessment were further discussed.

## Materials and Methods

### Cell growth and bacterial strains

*E. coli* K-12 ATCC 10798 was obtained from the American Type Culture Collection (Manassas, VA, USA). The streptomycin-resistant strain M14 was isolated from an *E. coli* K-12 ATCC 10798 population exposed to sub-MIC streptomycin with mixed pesticides after 500 generations.^14^ All other strains were generated by genetic manipulation of these two strains, as described below. Cells were grown in Luria–Bertani (LB) broth (10 g tryptone, 5 g yeast extract, and 10 g NaCl per liter).

### Genetic manipulation

The λ-red homologous recombination system was employed for the genetic manipulation of *E. coli*.^18^ The *rsmG* frameshift mutation (W150fs) was first introduced without leaving a selection marker. However, the markerless gene replacement workflow will not work for essential genes like *rpsE* and *rpsL*. Alternatively, a kanamycin resistance cassette (*Kan^R^*) was inserted downstream of *rpsL* and a chloramphenicol resistance cassette (*Cm^R^*) downstream of *rpsE*, without gene deletion. Retention of these selection markers in the genome did not affect streptomycin resistance (**Table S1**). All constructed strains, as well as primers and plasmids used are listed in **Table S2-S4**. A comprehensive list was compiled consisting of *rpsL* site mutations in streptomycin-resistant *E. coli* strains reported in the literature (**Table 1**). The corresponding site-specific mutants were constructed to examine their epistatic interactions with the other commonly found ribosomal mutation, *rsmG^W150fs^*.

**Table 1.** A list of *rpsL* point mutations investigated in this study.

| SNP<sup>a</sup> | Amino acid change | MIC of native isolates<sup>b</sup> reported in literature (mg/L) | MIC of *E. coli* mutant with the respective single *rpsL* mutation in this study<sup>c</sup> (mg/L) |
| --- | --- | --- | --- |
| C125T | P42L | > 1,200<sup>19</sup> | Lethal mutation<sup>d</sup> |
| A128C | K43T | > 10,000<sup>20</sup> | > 32,768 |
| A130G | K44E | > 1,200<sup>19</sup> | Lethal mutation |
| T221C | L74P | 25<sup>21</sup> | Lethal mutation |
| A230C | H77P | 25<sup>21</sup> | 64 |
| C256A | R86S | 100<sup>22</sup>; 25<sup>21</sup> | 64 |
| A263G | K88R | >10,000<sup>20</sup> | > 32,768 |
| C272T | P91L | > 10,000<sup>20</sup>; > 1,200<sup>19</sup> | Lethal mutation |
| G275A | G92D | > 10,000<sup>20</sup>; > 1,200<sup>19</sup> | Lethal mutation |

<sup>a</sup>Single nucleotide polymorphism (SNP) identified by Sanger sequencing of PCR products targeting *rpsL*.
<sup>b</sup>*E. coli* isolates were obtained in the lab by random mutagenesis or exposure to high-level streptomycin.
<sup>c</sup>Streptomycin MIC for the wild-type *E. coli* strain used in this study was 32 mg/L.
<sup>d</sup>Introduction of these *rpsL* point mutations into either the wild-type strain or the *rsmG*<sup>W150fs</sup> mutant was unsuccessful, due to lethality of those *rpsL* mutations, suggesting the presence of potential co-occurring unknown compensatory mutations.

### Growth curve measurement

Growth curves were measured in clear 96-well microplates. Archived wild-type and genetically manipulated mutants were revived and streaked on LB agar plates with respective selection markers (for mutants). A single colony was first picked up and inoculated into LB broth. After growing overnight at 37℃ and 180 rpm, cell culture was diluted to OD_600_ (optical density at 600 nm) = 0.1 with 1× phosphate buffer saline (PBS). Subsequently, 0.5 µL of the standardized cell suspension was added to 199.5 µL of LB medium. The 96-well microplate was placed in the BioTek Synergy H1 microplate reader (Agilent Technologies, Santa Clara, CA, USA) at 37℃ with orbital shaking, and OD_600_ values were measured every 15 min for 24 h. Five biological replicates were included. Area under the curve (AUC) was calculated using the following equation and used to represent the total cell growth:

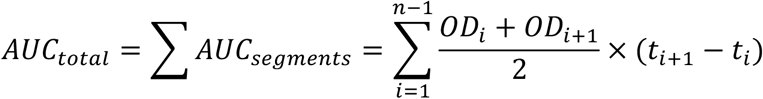

### Minimum inhibitory concentration (MIC) test

MIC was determined as the lowest antibiotic concentration that inhibited ≥ 90% of cell growth relative to the no-antibiotic control, as measured by OD_600_.^23^ MIC tests were performed in clear 96-well microplates. A single colony was first picked up and inoculated into LB broth. After growing overnight at 37℃ and 180 rpm, cell culture was diluted to OD_600_ = 0.1 with 1× PBS. Subsequently, 0.5 µL of the standardized cell suspension was added to 199.5 µL of LB medium supplemented with serial concentrations of streptomycin from 32 mg/L (MIC_0_) up to 32,768 mg/L (64× MIC_0_)). The 96-well microplate was then incubated at 37℃ for 20 h. After incubation, OD_600_ values were recorded by the BioTek Synergy H1 microplate reader. Three biological replicates were included.

### DNA extraction, whole-genome sequencing and variant calling

Genomic DNA of the streptomycin-resistant strain M14 was extracted using DNeasy Blood and Tissue Kits (Qiagen, Hilden, Germany), following the manufacturer’s instructions. The gDNA concentration was determined with a Qubit 4 Fluorometer (Thermo Fisher Scientific, Wilmington, DE, USA). Whole-genome sequencing was performed by SeqCenter (https://seqcenter.com, Pittsburgh, PA, USA) using an Illumina NovaSeq X Plus sequencer (2 × 150 bp). Raw data were trimmed by Trimmomatic v0.39,^24^ with a minimum length of 36 bp. Trimmed data were aligned against the genome of *E. coli* K-12 MG1655 downloaded from NCBI (accession: NC_000913) using Bowtie2 v2.5.1.^25^ SAMtools v1.2.2 and Picard v3.4.0 were applied to index and mark duplicates.^26^ Variant calling was performed by BCFtools v1.2.2 and SnpEff.^26, 27^ All identified mutations were manually checked in the genome, and genes of interest were confirmed by Sanger sequencing using the primers listed in **Table S1**. The sequenced genome has been deposited in NCBI Sequence Read Archive with the accession number PRJNA1332319.

## Results and Discussion

### Epistatic interactions among three ribosomal mutations conferred substantial increase in streptomycin resistance

Our previous study identified two ribosomal mutations (i.e., *rpsL^R86S^*and *rsmG^W150fs^*) co-occurring in *E. coli* mutants that evolved high-level streptomycin resistance (> 40-fold increase in MIC) under environmentally relevant selection pressures.^14, 28^ To disentangle the contribution of individual and combined mutations, we reconstructed each genotype in the wild-type, streptomycin-susceptible *E. coli* strain. Individual *rpsL^R86S^* or *rsmG^W150fs^*only led to twofold increase in MIC (**Figure 1A**). In contrast, their combination resulted in a 64-fold increase—far exceeding the simple additive effect of the two—thereby demonstrating an epistatic interaction between the two mutations (**Figure 1A**). To further validate these synergistic effects, we reverted the mutations to their wild-type alleles in the native resistant isolate M14, which evolved from the same wild-type strain under environmentally relevant exposures. As expected, reverting either *rpsL^R86S^*or *rsmG^W150fs^* caused a drastic drop in resistance (from 64-fold to 2-fold MIC_0_), and reverting both mutations fully restored streptomycin susceptibility (**Figure 1B**).

**Figure 1.**
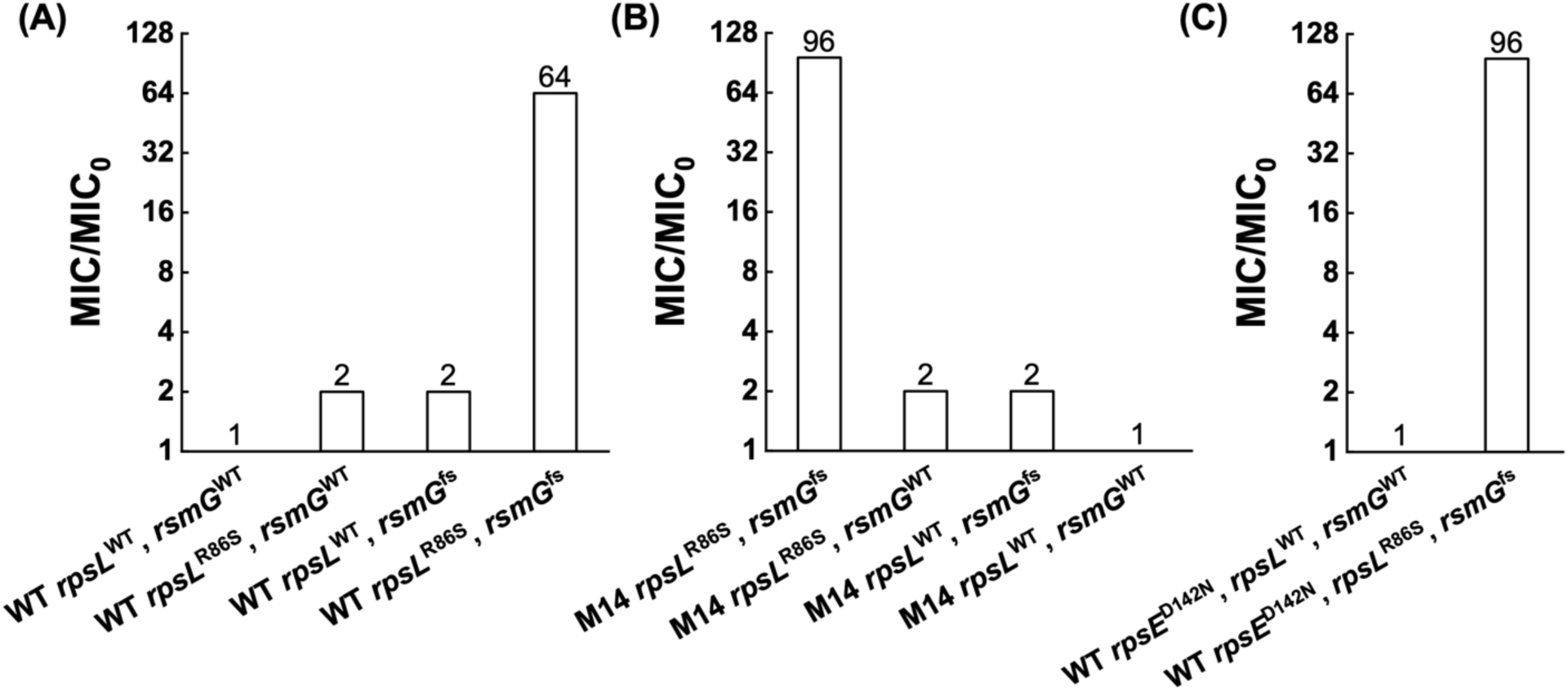
Effects of *rpsL*, *rsmG*, and *rpsE* mutations on streptomycin resistance. Minimum inhibitory concentration (MIC) fold changes relative to the wild type (WT, MIC_0_ = 32 mg/L) in different genetic backgrounds: (A) WT with *rpsL^R86S^* and *rsmG^W150fs^*mutations; (B) M14 with *rpsL^R86S^* and *rsmG^W150fs^*; (C) WT with *rpsE^D142N^* combined with *rpsL^R86S^* and *rsmG^W150fs^*. Exact MIC/MIC_0_ fold change values are shown above bars.

Notably, although introducing *rpsL^R86S^* and *rsmG^W150fs^*into the wild-type background reproduced the majority (67%) of the resistance observed in M14, the reconstructed strain still exhibited 33% lower resistance than that of M14 (64× vs. 96× MIC_0_). There seemed to be other mutations contributing to the overall resistance phenotype of M14, but the information was missing as only Sanger sequencing for *rpsL* and *rsmG* genes was done in the previous study. To test this, we further sequenced the genome of M14 and analyzed all missense mutations. Among them, we found another ribosomal mutation *rpsE^D142N^*. While introducing *rpsE^D142N^* alone into the wild type did not alter resistance, combining it with *rpsL^R86S^* and *rsmG^W150fs^* led to an additional increase of the resistance to the same level as that of M14 (**Figure 1C**).

Together, we demonstrated that it was the epistatic interactions among three ribosomal mutations, *rpsL^R86S^*, *rsmG^W150fs^*, and *rpsE^D142N^*, that conferred the high-level streptomycin resistance of M14 previously isolated from *E. coli* populations under environmentally relevant co-exposure to streptomycin and pesticides.^14^ The interaction between *rpsL^R86S^* and *rsmG^W150fs^* played a more significant role, with a small and incremental contribution by the third mutation *rpsE^D142N^*. These findings indicate the critical, but previously overlooked, role of epistatic interactions among co-evolved mutations in modulating high-level antibiotic resistance phenotypes.

### Epistatic interactions of *rsmG^W150fs^* with other *rpsL* mutations identified in streptomycin-resistant *E. coli* strains

In addition to *rpsL^R86S^*, several other *rpsL* point mutations have been reported in *E. coli* isolates exhibiting varying levels of streptomycin resistance (**Table 1**). However, the lack of whole-genome sequencing data of resistant *E. coli* strains obtained from random mutagenesis or antibiotic selection makes it unclear whether the resistance was solely caused by the reported *rpsL* mutation or by epistatic interactions with co-evolved mutations—potentially including *rsmG^W150fs^*, as demonstrated above. To address this knowledge gap, we investigated the interactions between *rsmG^W150fs^* and eight previously reported *rpsL* point mutations (**Table 1**). Three *rpsL* point mutations (H77P, K43N, and K88R) were successfully introduced into the wild-type strain, whereas the other five (P42L, K44E, L74P, P91L, and G92D) could not be obtained in either the wild-type or the *rsmG^W150fs^* mutant strain. Because *rpsL* encodes the essential ribosomal protein S12,^29^ these point mutations could be lethal or inhibitory to cell growth.

To rule out technical artifacts, we requested a commercial lab (GenScript Biotech, https://www.genscript.com) to introduce the *rpsL^P42L^* mutation into the wild type and the *rsmG^W150fs^*mutant using CRISPR-Cas9-based mutagenesis. Consistently, no viable mutants can be recovered. Thus, it most likely points to the lethality of those single *rpsL* mutations and the presence of unknown compensatory mutations in the native host, which were overlooked as only the *rpsL* gene was sequenced in those previous studies. Due to the lack of genome information, little further investigation could be done. This implies a critical role potentially played by multi-locus interactions in conferring a sustainable antibiotic resistance phenotype. It also underscores the need to identify all those interacting mutations as combined and more accurate antibiotic resistance biomarkers.

Among the successfully reconstructed *rpsL* point mutations, *rpsL^K43N^* and *rpsL^K88R^* are the two most frequently detected mutations, and their contribution to streptomycin resistance has been previously reported.^10^ Consistent with previous findings, both single mutants exhibited extremely high MIC (> 32,768 mg/L, > 1,024× MIC_0_), with detectable cell growth even at the highest streptomycin concentration tested. Introducing the *rsmG^W150fs^* mutation did not further increase the MIC, likely because resistance conferred by the *rpsL* mutation alone already exceeded the highest level tested. Thus, these *rpsL* alleles can independently confer extreme streptomycin resistance without requiring additional epistatic supports.

Meanwhile, we took a closer look at the effect of the additional *rsmG^W150fs^* mutation on growth fitness under typical streptomycin stresses. Growth curves were obtained under different streptomycin concentrations, and the area under the curve (AUC) was used to quantify relative fitness. Under streptomycin stress, the double mutants with both *rpsL* (K43N or K88R) and *rsmG^W150fs^* exhibited higher growth than the corresponding single *rpsL* mutant (**Figure 2** and **Figure S1**). Notably, the additional *rsmG^W150fs^* mutation also promoted a better fitness of the *rpsL^K43N^* mutant in the absence of streptomycin (**Figure 2A**). Therefore, although extremely high streptomycin resistance can be resulted by acquiring either of the two specific *rpsL* mutations, possessing another ribosomal mutation, *rsmG^W150fs^*, could provide a modest but measurable fitness advantage under both antibiotic selective and non-selective conditions.

**Figure 2.**
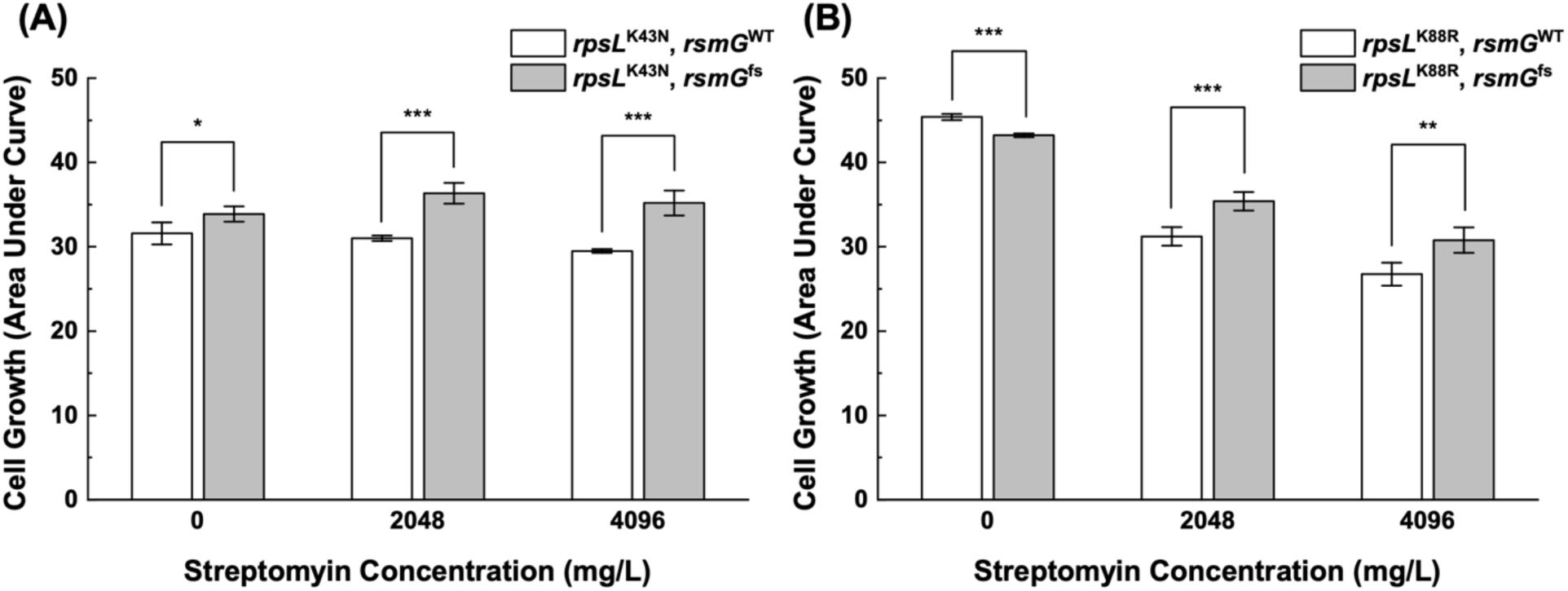
Growth of *rpsL* mutant strains with either *rsmG^WT^* or *rsmG^W150fs^*across increasing streptomycin concentrations. Cell growth was quantified as area under the curve (AUC). (A) *rpsL^K43N^*; (B) *rpsL^K88R^*. Data represent mean ± SD of three biological replicates. Asterisks denote the statistical significance of student *t*-test (*: *p* < 0.05; **: *p* < 0.01; ***: *p* < 0.001; n = 3).

Introducing *rpsL^H77P^* led to a mild increase in MIC (2 ×), similar to that observed in its native host (**Table 1**). When both *rpsL^H77P^* and *rsmG^W150fs^*were present, the MIC was increased by four times (**Figure 3A**), suggesting an additive effect different from the epistasis observed between *rpsL^R86S^* (the single mutation conferred mild resistance) and *rsmG^W150fs^*. It is worth noting that the *rpsL^H77P^* single mutant and the *rpsL^H77P^* + *rsmG^W150fs^* double mutant exhibited a remarkable fitness cost with a more significant decrease in cell growth for the double mutant (**Figure 3B**). The impaired cell growth could compromise the resistance phenotype of those mutants.

**Figure 3.**
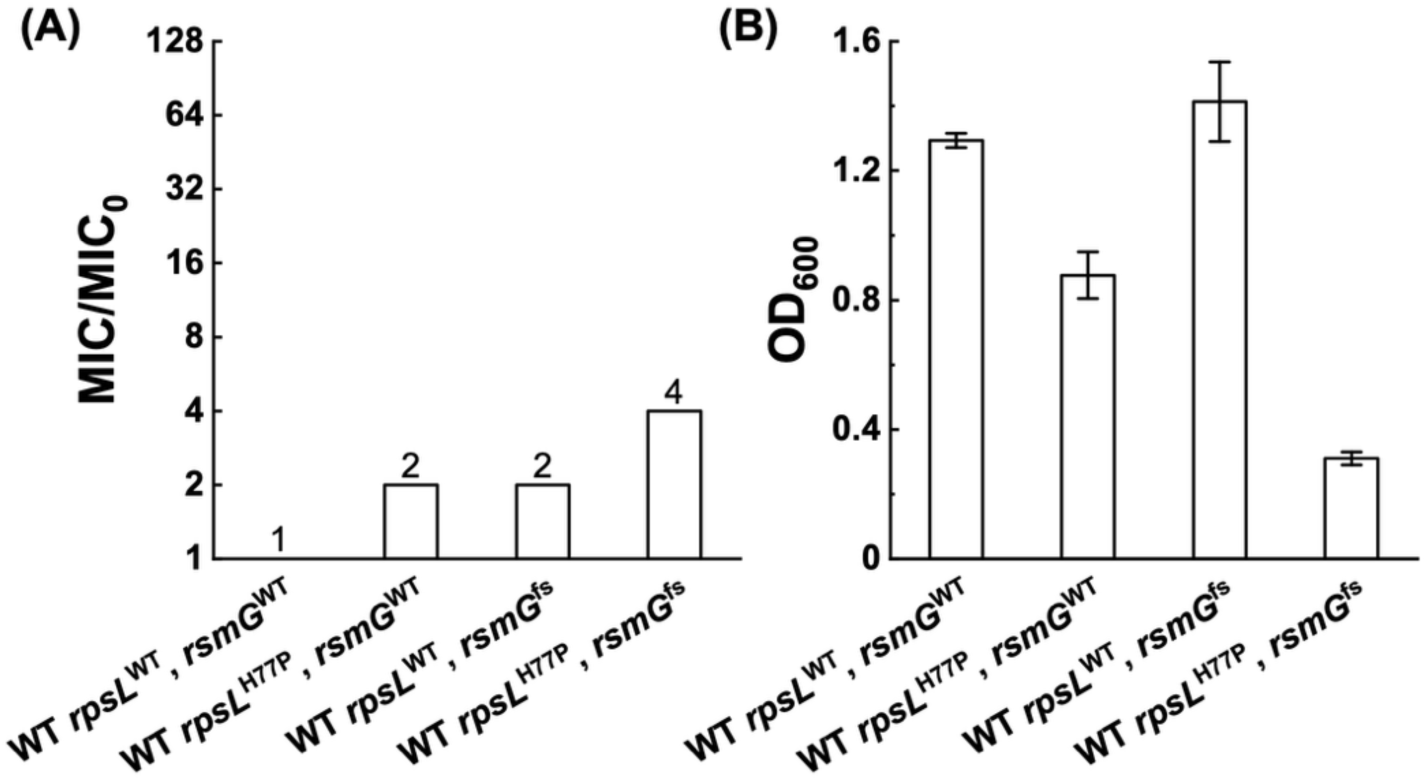
Effects of *rpsL*^H77P^ mutations on. (A) streptomycin resistance and (B) cell growth with either *rsmG^WT^* or *rsmG^W150fs^* in streptomycin-free LB medium. Exact MIC/MIC_0_ fold change values are shown above bars. OD_600_ was measured after 20 h. Data represent mean ± SD of three biological replicates.

Collectively, other previously reported *rpsL* mutations did not show epistatic interactions with *rsmG*^W150fs^, although having *rsmG*^W150fs^ added some growth benefit for the *rpsL^K43N^*or *rpsL^K88R^* mutants. One possible reason is that the epistatic interactions were specific to *rpsL* mutation sites. It can also be explained by the natural selection process, where only sustainable mutations could be selected and remain under the antibiotic selection pressure. Lethal mutations and mutations with substantial fitness cost may not sustain, and it was the case for several *rpsL* single mutants and the *rpsL^H77P^*+ *rsmG* double mutant in our study. Our previous studies revealed that *rsmG* loss-of-function mutations (e.g., *rsmG^W150fs^*) co-evolved with one specific *rpsL* site mutation *rpsL^R86S^*.^14,28^ In the same study,^28^ *rpsL^K88R^* and *rpsL^K43N^* mutations were detected without any accompanying *rsmG* mutations in *E. coli* populations under the co-exposure of streptomycin and pesticides. Since the *rpsL^K43N^* or *rpsL^K88R^*single mutation already showed extremely high resistance with no significant sacrifice on growth, the growth benefit provided by the additional *rsmG* mutation seemed to be marginal and not sufficient to have the double mutant to stand out or even emerge during the typical lab evolution process. However, whether *rsmG* co-evolved with other *rpsL* mutations in **Table 1** remains elusive due to unavailable genome information of the corresponding resistant mutants. It again emphasizes the need for whole-genome analysis of antibiotic-resistant bacteria to identify co-evolved mutations (e.g., *E. coli* strains with lethal *rpsL* single mutations) that could compensate fitness cost and confer strong antibiotic resistance via epistatic interactions.

## Environmental Implications

The findings of this study fill a large knowledge gap and advance the fundamental understanding of antibiotic resistance mechanisms mediated by target-altering mutations, thereby can benefit antibiotic resistance surveillance. In addition to the horizontal transfer of antibiotic resistance genes, *de novo* mutations causing target alterations represent another major route of resistance development to many classes of antibiotics. Such mutations can reduce antibiotic binding affinity, thereby conferring resistance. For example, mutations of genes encoding proteins involved in protein and DNA synthesis have been identified in aminoglycoside and fluoroquinolone-resistant bacteria, respectively. In natural and agricultural settings, bacteria are frequently exposed to sub-inhibitory concentrations of antibiotics along with other chemical stressors such as pesticides.^13^ The non-lethal exposure could facilitate more diverse *de novo* mutations across both antibiotic target and non-target genes. Here, we demonstrated that the high-level streptomycin resistance observed in resistant isolates under non-lethal exposures was attributed to epistatic interactions among ribosomal mutations (e.g., *rpsL* and *rsmG*), each of which alone only conferred mild resistance. This suggests that the contribution of such epistatic interactions in resistance development in a broader environmental context could have been largely overlooked, particularly because most previous studies focused on individual genetic mutations.

Under environmentally relevant exposures, the same epistasis-mediated resistance mechanisms may be found among mutations of genes encoding a wide range of antibiotic targets, including DNA gyrases and topoisomerases (GyrAB and ParCE) for fluoroquinolones and RNA polymerases (e.g., RpoB) for rifampicin.^15, 30^ Therefore, to capture such interactions, future studies should employ whole genome analysis rather than single-gene sequencing to get holistic mutation profiles of antibiotic-resistant mutants/populations, and even environmental microbial communities. This would allow us to comprehensively interrogate potential epistatic interactions among co-evolved mutations for an extended range of antibiotics. Whole-genome/population sequencing has been achieved for large sample sizes at an affordable cost. The advancement of long-length and deep sequencing and the decreasing sequencing cost could help realize the detection of genetic mutations in complex environmental samples. Thus, validated individual and epistasis-based resistance mutations can be incorporated with resistomes analysis to curate sequencing-based surveillance and promote a more accurate antibiotic resistance monitoring and risk assessment.

## Acknowledgements

This study was supported by the OASIS Internal Funding Award at University of California, Riverside and the National Science Foundation (Award No. 2045658).

## Notes

### Competing Interest Statement

The authors have declared no competing interest.

## References

(1) Larsson, D. G. J.; Flach, C. F. Antibiotic resistance in the environment. Nat Rev Microbiol 2022, 20 (5), 257–269.

(2) Centers for Disease Control and Prevention. Antimicrobial Resistance Threats in the United States, 2021–2022; U.S. Department of Health and Human Services, Atlanta, GA, 2025. https://www.cdc.gov/antimicrobial-resistance/data-research/threats/update-2022.html.

(3) European Centre for Disease Prevention and Control (ECDC); WHO Regional Office for Europe. Surveillance of antimicrobial resistance in Europe, 2023 data: executive summary; European Centre for Disease Prevention and Control, Stockholm, Sweden, 2024. https://www.ecdc.europa.eu/en/publications-data/surveillance-antimicrobial-resistance-europe-2023-data-executive-summary.

(4) Crofts, T. S.; Gasparrini, A. J.; Dantas, G. Next-generation approaches to understand and combat the antibiotic resistome. Nat Rev Microbiol 2017, 15 (7), 422–434.

(5) Zhu, C.; Wu, L.; Ning, D.; Tian, R.; Gao, S.; Zhang, B.; Zhao, J.; Zhang, Y.; Xiao, N.; Wang, Y.;, et al. Global diversity and distribution of antibiotic resistance genes in human wastewater treatment systems. Nature Communications 2025, 16 (1), 4006.

(6) McManus, P. S.; Stockwell, V. O.; Sundin, G. W.; Jones, A. L. Antibiotic use in plant agriculture. Annu Rev Phytopathol 2002, 40, 443–465.

(7) Stockwell, V. O.; Duffy, B. Use of antibiotics in plant agriculture. Rev Sci Tech 2012, 31 (1), 199–210.

(8) Davies, J.; Davies, D. Origins and evolution of antibiotic resistance. Microbiol Mol Biol Rev 2010, 74 (3), 417–433.

(9) Boolchandani, M.; D’Souza, A. W.; Dantas, G. Sequencing-based methods and resources to study antimicrobial resistance. Nat Rev Genet 2019, 20 (6), 356–370.

(10) Pelchovich, G.; Schreiber, R.; Zhuravlev, A.; Gophna, U. The contribution of common rpsL mutations in Escherichia coli to sensitivity to ribosome targeting antibiotics. Int J Med Microbiol 2013, 303 (8), 558–562.

(11) Okamoto, S.; Tamaru, A.; Nakajima, C.; Nishimura, K.; Tanaka, Y.; Tokuyama, S.; Suzuki, Y.; Ochi, K. Loss of a conserved 7-methylguanosine modification in 16S rRNA confers low-level streptomycin resistance in bacteria. Mol Microbiol 2007, 63 (4), 1096–1106.

(12) Wong, S. Y.; Lee, J. S.; Kwak, H. K.; Via, L. E.; Boshoff, H. I.; Barry, C. E., 3rd. Mutations in gidB confer low-level streptomycin resistance in Mycobacterium tuberculosis. Antimicrob Agents Chemother 2011, 55 (6), 2515–2522.

(13) Luo, Y.; Guo, W.; Ngo, H. H.; Nghiem, L. D.; Hai, F. I.; Zhang, J.; Liang, S.; Wang, X. C. A review on the occurrence of micropollutants in the aquatic environment and their fate and removal during wastewater treatment. Sci Total Environ 2014, 473-474, 619–641.

(14) Xing, Y.; Kang, X.; Zhang, S.; Men, Y. Specific phenotypic, genomic, and fitness evolutionary trajectories toward streptomycin resistance induced by pesticide co-stressors in Escherichia coli. ISME Commun 2021, 1 (1), 39.

(15) Xing, Y.; Wu, S.; Men, Y. Exposure to Environmental Levels of Pesticides Stimulates and Diversifies Evolution in Escherichia coli toward Higher Antibiotic Resistance. Environ Sci Technol 2020, 54 (14), 8770–8778.

(16) Howarth, R. E.; Pattillo, C. M.; Griffitts, J. S.; Calvopina-Chavez, D. G. Three genes controlling streptomycin susceptibility in Agrobacterium fabrum. J Bacteriol 2023, 205 (9), e0016523.

(17) Tanaka, Y.; Komatsu, M.; Okamoto, S.; Tokuyama, S.; Kaji, A.; Ikeda, H.; Ochi, K. Antibiotic overproduction by rpsL and rsmG mutants of various actinomycetes. Appl Environ Microbiol 2009, 75 (14), 4919–4922.

(18) Datsenko, K. A.; Wanner, B. L. One-step inactivation of chromosomal genes in Escherichia coli K-12 using PCR products. Proc Natl Acad Sci U S A 2000, 97 (12), 6640–6645.

(19) Timms, A. R.; Steingrimsdottir, H.; Lehmann, A. R.; Bridges, B. A. Mutant sequences in the rpsL gene of Escherichia coli B/r: mechanistic implications for spontaneous and ultraviolet light mutagenesis. Mol Gen Genet 1992, 232 (1), 89–96.

(20) Chumpolkulwong, N.; Hori-Takemoto, C.; Hosaka, T.; Inaoka, T.; Kigawa, T.; Shirouzu, M.; Ochi, K.; Yokoyama, S. Effects of Escherichia coli ribosomal protein S12 mutations on cell-free protein synthesis. Eur J Biochem 2004, 271 (6), 1127–1134.

(21) Agarwal, D.; Gregory, S. T.; O’Connor, M. Error-prone and error-restrictive mutations affecting ribosomal protein S12. J Mol Biol 2011, 410 (1), 1–9.

(22) Edgar, R.; Friedman, N.; Molshanski-Mor, S.; Qimron, U. Reversing bacterial resistance to antibiotics by phage-mediated delivery of dominant sensitive genes. Appl Environ Microbiol 2012, 78 (3), 744–751.

(23) Kohanski, M. A.; DePristo, M. A.; Collins, J. J. Sublethal antibiotic treatment leads to multidrug resistance via radical-induced mutagenesis. Mol Cell 2010, 37 (3), 311–320.

(24) Bolger, A. M.; Lohse, M.; Usadel, B. Trimmomatic: a flexible trimmer for Illumina sequence data. Bioinformatics 2014, 30 (15), 2114–2120.

(25) Langmead, B.; Salzberg, S. L. Fast gapped-read alignment with Bowtie 2. Nat Methods 2012, 9 (4), 357–359.

(26) Danecek, P.; Bonfield, J. K.; Liddle, J.; Marshall, J.; Ohan, V.; Pollard, M. O.; Whitwham, A.; Keane, T.; McCarthy, S. A.; Davies, R. M.;, et al. Twelve years of SAMtools and BCFtools. Gigascience 2021, 10 (2).

(27) Cingolani, P.; Platts, A.; Wang le, L.; Coon, M.; Nguyen, T.; Wang, L.; Land, S. J.; Lu, X.; Ruden, D. M. A program for annotating and predicting the effects of single nucleotide polymorphisms, SnpEff: SNPs in the genome of Drosophila melanogaster strain w1118; iso-2; iso-3. Fly (Austin) 2012, 6 (2), 80–92.

(28) Xing, Y.; Herrera, D.; Zhang, S.; Kang, X.; Men, Y. Site-specific target-modification mutations exclusively induced by the coexposure to low levels of pesticides and streptomycin caused strong streptomycin resistance in clinically relevant Escherichia coli. Journal of Hazardous Materials Advances 2022, 7, 100141.

(29) Sharma, D.; Cukras, A. R.; Rogers, E. J.; Southworth, D. R.; Green, R. Mutational analysis of S12 protein and implications for the accuracy of decoding by the ribosome. J Mol Biol 2007, 374 (4), 1065–1076.

(30) Lambert, P. A. Bacterial resistance to antibiotics: Modified target sites. Advanced Drug Delivery Reviews 2005, 57 (10), 1471–1485.

