## Supplemental Table S1-S4; Figure S1 for "Ribosomal Epistatic Interactions Drive the Emergence of High-Level Streptomycin Resistance in *Escherichia coli*"

**Table S1. Influence of introducing *KanR/CmR* on streptomycin resistance.**

| #Strain | Description | MIC (mg/L) |
|---|---|---|
| FX-500 | wild-type *E. coli* K-12 ATCC No.10798 strain, rpsL(WT)::rsmG(WT) | 32 |
| FX-526 | wild-type strain, rpsL(WT)::KanR::rsmG(WT) | 32 |
| FX-553 | wild-type strain, rpsE(WT)::CmR::rpsL(WT)::KanR::rsmG(WT) | 32 |

**Table S2. Primers used in this study.**

| Primer Name | Primer Seuqence (5' - 3') |
|---|---|
| rpsL_42_F | CTTAAAAAACCGAACTCCGCGC |
| rpsL_42_R | GCGCGGAGTTCGGTTTTTTAAG |
| rpsL_43_F | CCTACAAAACCGAACTCCGCGC |
| rpsL_43_R | GCGCGGAGTTCGGTTTTGTAGG |
| rpsL_44_F | CCTAAAGAACCGAACTCCGCGC |
| rpsL_44_R | GCGCGGAGTTCGGTTCTTTAGG |
| rpsL_74_F | GTGGTGAAGGTCACAACCCGCA |
| rpsL_74_R | TGCGGGTTGTGACCTTCACCAC |
| rpsL_77_F | AGGAGCCCTCCGTGATCCTGA |
| rpsL_77_R | TCAGGATCACGGAGGGCTCCT |
| rpsL_88_F | CGTGTTAGAGACCTCCCGGGT |
| rpsL_88_R | ACCCGGGAGGTCTCTAACACG |
| rpsL_91_F | CGTGTTAAAGACCTCCTGGGT |
| rpsL_91_R | ACCCAGGAGGTCTTTAACACG |
| rpsL_92_F | CGTGTTAAAGACCTCCCGGAT |
| rpsL_92_R | ATCCGGGAGGTCTTTAACACG |
| rpsL_F | ATGGCAACAGTTAACCAGCTGG |
| rpsL_R | AATCCATCTTGTTCAATCATGCG<br>TTAAGCCTTAGGACGCTTCACGC |
| in_rpsL_KanR_F | GCGTGAAGCGTCCTAAGGCTTAACGC<br>ATGATTGAACAAGATGGATT |
| in_rpsL_KanR_R | TTAGTTTGACATTTAAGTTAAAACGTTTGGCCTTACTTAACGGAGAACCA<br>TCAGAAGAACTCGTCAAG |
| rsmG_F | GTGCTCAACAAACTCTCCTTACTGC |
| rsmG_R | AATCCATCTTGTTCAATCATGCG<br>TTAAATTTTATTTGCTTTAATCACCACCAG |
| rsmG_KanR_F | GTGGTGATTAAAGCAAATAAAATTTAA<br>TTGTGTAGGCTGGAGCTGCTTC |
| rsmG_KanR_R | TGTTGTTAACAGTCTAACCGGTCAATTTTTTATGATTTTTTTGATAAAAA<br>CATATGAATATCCTCCTTAGTTCCTATTCC |
| rpsE_F | ATGGCTCACATCGAAAAACAAGCTG |
| rpsE_R | CTAAGGAGGATATTCATATG<br>TTATTTCCCCAGAATTTCTTCAACGGATTT |
| in_rpsE_CmR_F | CATATGAATATCCTCCTTAGTTCCTATTCC |
| in_rpsE_CmR_R | CGACCGATTGCACTGCGGGTTTGAGTAATTTTAATAGTCTTTGCCATGGT<br>TTGTGTAGGCTGGAGCTGCTTC |

**Table S3. Plasmids used in this study.**

| Plasmid Name | Source |
|---|---|
| pKD3 | Addgene[1] |
| pKD4 | Addgene[1] |
| pKM208 | Addgene[2] |
| pBAD-Flp | Addgene[3] |

**Table S4.** **Bacterial strains used in this study.**

| #Strain | Description | Reference |
|---|---|---|
| FX-500 | wild-type *E. coli* K-12 ATCC No.10798 strain | ATCC |
| FX-501 | isolate M14 | [4] |
| FX-518 | wild-type strain, rpsL(86)::KanR::rsmG(WT) | This study |
| FX-520 | wild-type strain, ΔrsmG::rsmG(fs) | This study |
| FX-526 | wild-type strain, rpsL(WT)::KanR::rsmG(WT) | This study |
| FX-527 | wild-type strain, rpsL(WT)::KanR::rsmG(fs) | This study |
| FX-528 | wild-type strain, rpsL(86)::KanR::rsmG(fs) | This study |
| FX-529 | isolate M14, rpsL(WT)::KanR::rsmG(WT) | This study |
| FX-530 | isolate M14, rpsL(86)::KanR::rsmG(WT) | This study |
| FX-531 | isolate M14, rpsL(WT)::KanR::rsmG(fs) | This study |
| FX-532 | isolate M14, rpsL(86)::KanR::rsmG(fs) | This study |
| FX-533 | wild-type strain, rpsL(43)::KanR::rsmG(WT) | This study |
| FX-534 | wild-type strain, rpsL(43)::KanR::rsmG(fs) | This study |
| FX-538 | wild-type strain, rpsL(88)::KanR::rsmG(WT) | This study |
| FX-539 | wild-type strain, rpsL(88)::KanR::rsmG(fs) | This study |
| FX-544 | wild-type strain, rpsL(77)::KanR::rsmG(WT) | This study |
| FX-545 | wild-type strain, rpsL(77)::KanR::rsmG(fs) | This study |
| FX-553 | wild-type strain, rpsE(WT)::CmR::rpsL(WT)::KanR::rsmG(WT) | This study |
| FX-554 | wild-type strain, rpsE(142)::CmR::rpsL(WT)::KanR::rsmG(WT) | This study |
| FX-555 | wild-type strain, rpsE(WT)::CmR::rpsL(86)::KanR::rsmG(fs) | This study |
| FX-556 | wild-type strain, rpsE(142)::CmR::rpsL(86)::KanR::rsmG(fs) | This study |

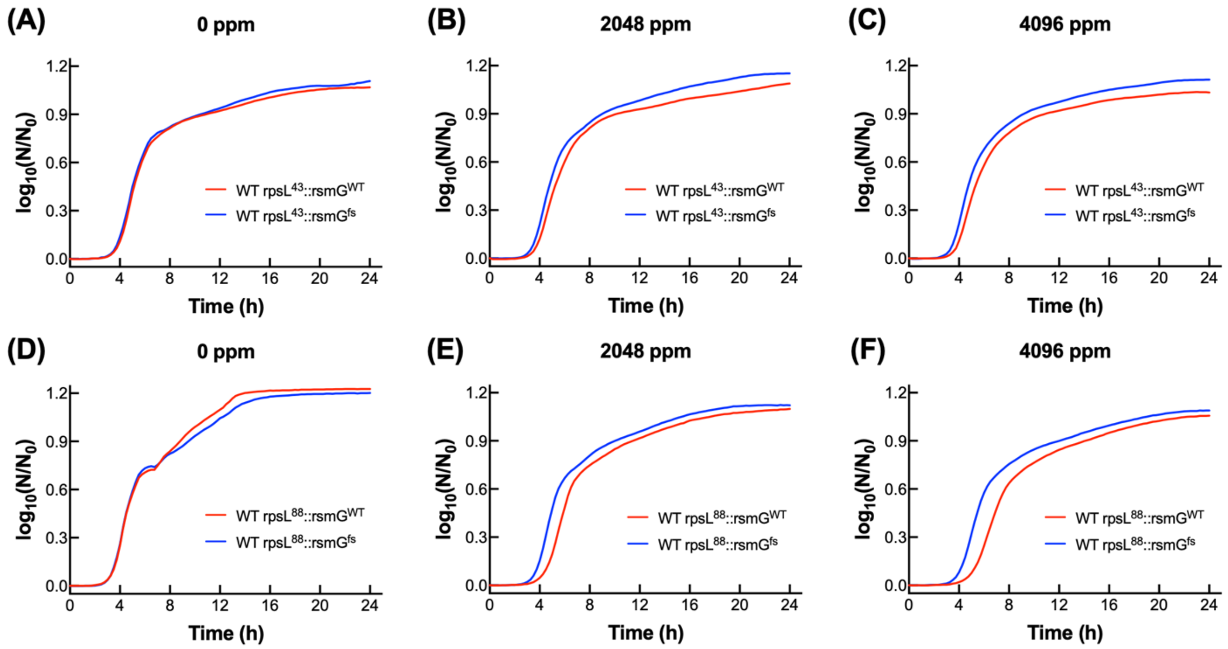


**Figure S1. Growth curve of *rpsL* mutant strains with either *rsmG*$^{WT}$ or *rsmG*$^{W150fs}$ across increasing streptomycin concentrations**. $OD_{600}$ (optical density at 600 nm) was measured every 15 min and log transformed. (A-C) *rpsL*$^{K43N}$; (D-F) *rpsL*$^{K88R}$. Three biological replicates were included for each group.

### References

(1) Datsenko, K. A.; Wanner, B. L. One-step inactivation of chromosomal genes in Escherichia coli K-12 using PCR products. *Proc Natl Acad Sci U S A* **2000**, *97* (12), 6640-6645. DOI: 10.1073/pnas.120163297 From NLM Medline.

(2) Murphy, K. C.; Campellone, K. G. Lambda Red-mediated recombinogenic engineering of enterohemorrhagic and enteropathogenic E. coli. *BMC Mol Biol* **2003**, *4*, 11. DOI: 10.1186/1471-2199-4-11 From NLM Medline.

(3) Dalia, T. N.; Chlebek, J. L.; Dalia, A. B. A modular chromosomally integrated toolkit for ectopic gene expression in Vibrio cholerae. *Sci Rep* **2020**, *10* (1), 15398. DOI: 10.1038/s41598-020-72387-8 From NLM Medline.

(4) Xing, Y.; Kang, X.; Zhang, S.; Men, Y. Specific phenotypic, genomic, and fitness evolutionary trajectories toward streptomycin resistance induced by pesticide co-stressors in Escherichia coli. *ISME Commun* **2021**, *1* (1), 39. DOI: 10.1038/s43705-021-00041-z From NLM PubMed-not-MEDLINE.
